# Five decades of data yield no support for adaptive biasing of offspring sex ratio in wild baboons (*Papio cynocephalus*)

**DOI:** 10.1101/2022.08.28.505562

**Authors:** Matthew N Zipple, Elizabeth A Archie, Jenny Tung, Raphael S. Mututua, J. Kinyua Warutere, I. Long’ida Siodi, Jeanne Altmann, Susan C Alberts

## Abstract

Over the past 50 years, a wealth of testable, often conflicting, hypotheses have been generated about the evolution of offspring sex ratio manipulation by mothers. Several of these hypotheses have received support in studies of invertebrates and some vertebrate taxa. However, their success in explaining sex ratios in mammalian taxa, and especially in primates, has been mixed. Here, we assess the predictions of four different hypotheses about the evolution of biased offspring sex ratios in the well-studied baboons of the Amboseli basin in Kenya: the Trivers-Willard, female rank enhancement, local resource competition, and local resource enhancement hypotheses. Using the largest sample size ever analyzed in a primate population (n = 1372 offspring), we test the predictions of each hypothesis. Overall, we find no support for adaptive biasing of sex ratios. Offspring sex is not consistently related to maternal dominance rank or biased towards the dispersing sex, nor it is predicted by group size, population growth rates, or their interaction with maternal rank. Because our sample size confers power to detect even subtle biases in sex ratio, including modulation by environmental heterogeneity, these results suggest that adaptive biasing of offspring sex does not occur in this population.

## Introduction

A wide range of evolutionary theories have been proposed to explain why mammalian mothers might adaptively bias the sex ratios of their offspring. These theories focus on two main types of maternal benefits to producing offspring of the “right” sex. First, by biasing offspring production to the optimal sex, mothers may be able to maximize their offspring’s lifetime reproductive success (the Trivers-Willard and female rank enhancement hypotheses). Second, biased sex ratios may help optimize the competitive environment for mothers themselves (the local resource competition and local resource enhancement hypotheses). All of these hypotheses make distinct, testable predictions about the influence of maternal condition, the competitive environment, and a species’ natural history on offspring sex ratio. However, while some hypotheses regarding adaptive sex ratio bias have been supported in invertebrates and birds (West & Sheldon 2002), their record in mammals, especially in primates, remains mixed (Brown & Silk 2002, Silk et al. 2005). Consequently, the importance—and even existence—of maternally driven sex ratio bias in primates remains in dispute (Brown 2001, Brown & Silk 2002, Silk et al. 2005).

### Maximizing offspring fitness: the Trivers-Willard and female rank enhancement hypotheses

The first way in which mothers might benefit from controlling their offspring’s sex is by increasing the reproductive success of their offspring. In many species, reproductive success depends on an individual’s physical condition, and variation in reproductive success differs between the sexes (sex-specific reproductive skew (Hauber & Lacey 2005)). As a result, a mother who can produce especially robust offspring should produce offspring of the sex with greater variance in reproductive success (usually males, but this pattern is reversed in singular-breeding cooperative societies, (Hauber & Lacey 2005)). In contrast, a mother who will produce offspring in poor condition should produce offspring of the sex with lower variance in reproductive success.

Trivers and Willard (1973) made two assumptions that led to strong predictions about the evolution of offspring sex ratios. They argued that if (1) maternal condition during the period of maternal investment affects offspring condition at the end of the period of maternal investment, and (2) sons differentially benefit from this increase in physical condition in terms of lifetime reproductive success as compared to daughters (due to sex-specific reproductive skew), then top-condition mothers should be selected to produce more sons compared to mothers in poor condition. These sons would then benefit from their mother’s condition, develop into high-quality adult males, and achieve greater reproductive success than if they had grown into high-quality adult females (Trivers and Willard, 1973).

Among nonhuman primates, the Trivers-Willard hypothesis was first evaluated in the wild yellow baboons (*Papio cynocephalus*) of the Amboseli ecosystem of Kenya, where maternal rank was used as a proxy for maternal condition (rank predicts a wide range of condition-related traits in this population, Levy and Zipple et al. 2020). In this setting, the Trivers-Willard hypothesis predicts that high-ranking mothers should produce relatively many sons, while low-ranking mothers should produce relatively many daughters (Table 1). Two studies from the Amboseli population (Altmann 1980, Altmann et al. 1988) identified a strong relationship between maternal rank and secondary sex ratio, *but in the opposite direction as originally predicted by Trivers and Willard*. Indeed, analyses of sex ratios of offspring born in Amboseli over a seven-year period (1971-1978; Altmann 1980) or an expanded ten-year period (1971-1981; Altmann et al. 1988) showed that high-ranking females were much more likely to give birth to daughters as compared to low-ranking females, and that this phenomenon was predicted by a continuous measure of rank (i.e., not just a binary metric of high-vs low-ranking females).

**Table 1.**
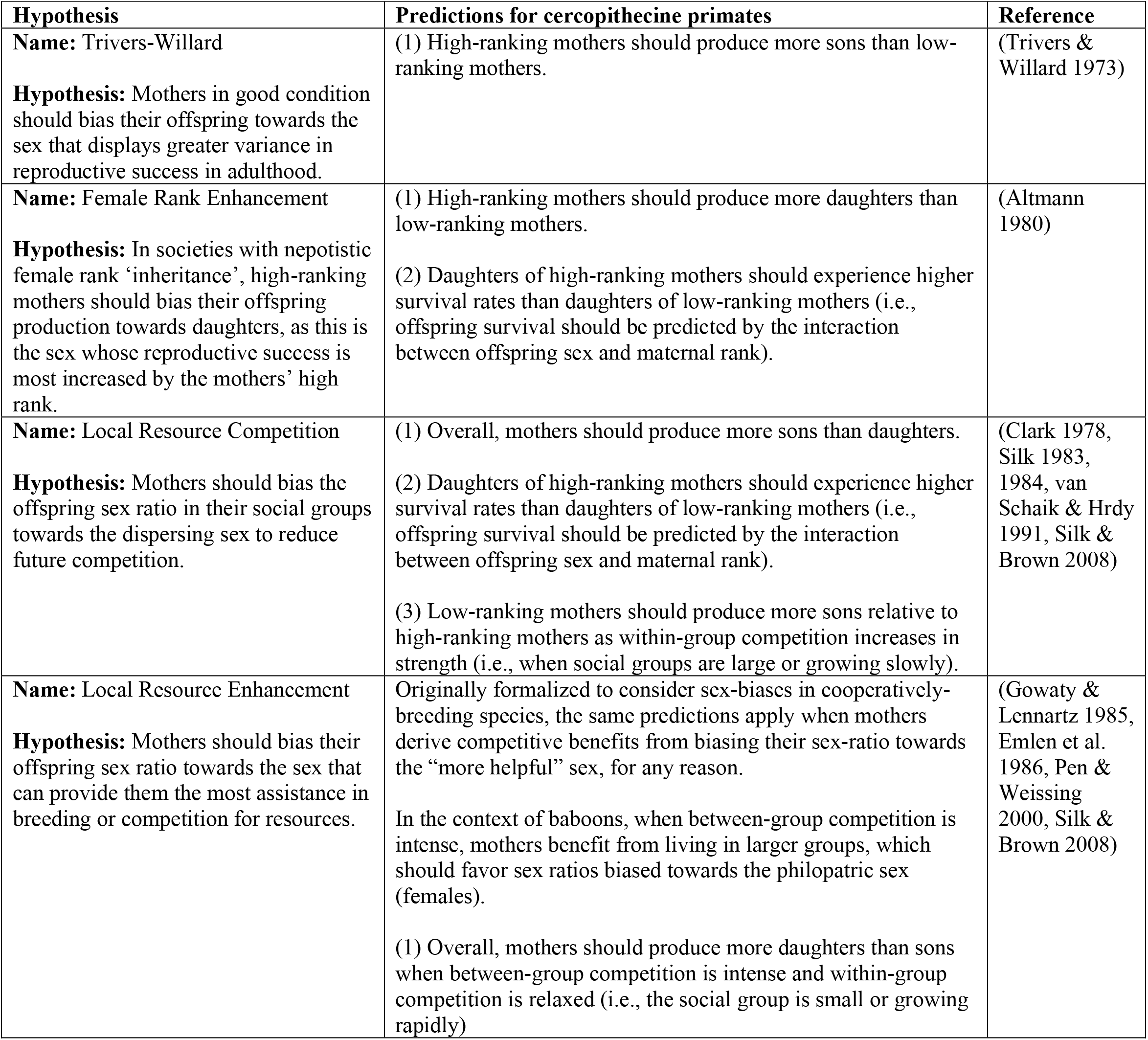
Hypotheses and predictions about adaptive biasing of offspring sex ratios, as they pertain to species with male-biased dispersal and matrilineal rank inheritance.

The results from Amboseli gave rise to an alternative hypothesis, which we term here the “female rank enhancement hypothesis.” This hypothesis reformulates the Trivers-Willard hypothesis to better fit the life history of mammals that exhibit matrilineal rank inheritance. Specifically, Altmann (1980) suggested that high-ranking females should bias their offspring sex ratio towards the sex whose reproductive success is most improved by the females’ high rank. Because cercopithecine females tend to inherit the rank of their mother in adulthood, while sons do not, Altmann posited that the first assumption of the Trivers-Willard hypothesis—that maternal condition during the period of investment shapes later offspring competitive ability— only held for female offspring. In the case of baboons, a high-ranking daughter would grow into a high-ranking adult and enjoy the fitness benefits that her high rank conferred (e.g. increased offspring survival and shorter interbirth intervals, (Silk et al. 2003, Gesquiere et al. 2018, Zipple et al. 2019), while a son born to a high-ranking mother would be no better off in the long run than a son born to a low-ranking mother. The female rank enhancement hypothesis therefore predicts that high-ranking baboon mothers should produce relatively many daughters, while low-ranking mothers should produce relatively many sons (Table 1). Female rank inheritance is common among cercopithecine primates, as well as some other taxa (e.g. spotted hyenas, (Strauss et al. 2020)), highlighting the potential generalizability of this hypothesis.

The results from Amboseli were followed by many similar analyses across at least 15 species of primates, testing the alternative predictions of the Trivers-Willard and female rank enhancement hypotheses (Reviewed in (Brown 2001)). Some of these studies were consistent with the female rank enhancement hypothesis (e.g. bonnet macaques: (Silk 1988); rhesus macaques: (Nevison et al. 1996)), while others found no effect of maternal rank on offspring sex ratio (e.g. yellow baboons, (Rhine et al. 1992); Toque macaques: (Dittus 1998); Japanese macaques: (Koyama et al. 1992), vervet monkeys: (Cheney et al. 1988)), and still others found an effect in the opposite direction, consistent with the Trivers-Willard hypothesis (i.e. high-ranking mothers had more sons than low-ranking mothers, e.g. rhesus macaques: (Meikle et al. 1984); Barbary macaques: (Paul & Kuester 1990); spider monkeys: (McFarland Symington 1987)).

A meta-analysis by Brown and Silk (2002) found that results from these and other studies of maternal rank and offspring sex in non-human primates did not collectively deviate from the expected null distribution of effect sizes, after controlling for the sample size of offspring included in each study. The meta-analysis supported neither the Trivers-Willard nor the female rank enhancement hypotheses, leaving unclear whether non-human primate mothers are capable of adjusting their offspring sex ratio based on social rank or other aspects of the environment.

However, studies with larger sample sizes also covered longer time periods and were likely affected by greater environmental heterogeneity than smaller studies (Brown & Silk 2002). Increased environmental heterogeneity may have made it more difficult to detect true sex biases if some environments favor such a bias while others do not. It therefore remains possible that mothers benefit from adjusting their offspring sex ratio as a function of rank in some contexts but not others. For example, Kruuk et. al. (1999) presented evidence that dominant red deer (*Cervus elaphus*) mothers bias their offspring sex ratios towards sons during periods of low population density, but that this effect disappears when density and resource competition is high. Failing to account for such heterogeneity could cause researchers to miss a real effect of maternal rank on offspring sex.

### Optimizing the competitive environment: the local resource competition and local resource enhancement hypotheses

The second way in which mothers could benefit from biasing the sex ratio of their offspring is by optimizing the competitive environment that they (the mothers and their offspring) experience. Clark (1978) argued that, in species that exhibit sex-biased dispersal—such that members of one sex generally disperse while members of the other sex do not—offspring sex determines whether mothers and offspring co-reside, cooperate, and compete in adulthood. For example, male baboons disperse while female baboons are philopatric, which results in female baboons co-residing with their adult daughters, but not their adult sons.

The local resource competition hypothesis argues that, when sons disperse and daughters are philopatric, females should benefit by limiting the production of daughters (both their own and other females’), thereby limiting the number of competitors in their immediate social group (Clark 1978). As a result, populations of female-philopatric species should display an overall bias towards sons—a bias that Clark first observed in greater galagos (*Galago crassicaudatus*), and a prediction that is supported in primates generally (Silk & Brown 2008), but not in baboons (Silk et al. 2005).

Silk (1983, 1984) extended the local resource competition hypothesis, arguing that females should (i) attempt to limit the survival of unrelated immature females and (ii) facultatively adjust their own offspring sex ratios depending on their competitive ability (i.e., their social rank). Thus, low-ranking females should show an especially strong bias towards sons, relative to high-ranking females (who might not bias towards sons at all, Silk 1983). Silk’s formulation of the local resource competition hypothesis dovetails with the female rank enhancement hypothesis in this prediction (Silk 1983,1984).

van Schaik and Hrdy (1991) further argued that the facultative sex ratio adjustment posited by Silk (1983, 1984) should depend on the intensity of resource competition, such that the relationship between maternal rank and offspring sex should intensify as competition for resources intensifies and population growth rate declines. Thus, the local resource competition hypothesis predicts that (1) at the population level, offspring sex ratios should be biased towards the dispersing sex, (2) low-ranking females should produce more sons than high-ranking females (consistent with the female rank enhancement hypothesis), (3) this rank-related sex bias should be especially apparent during periods of intense competition, and (4) low-ranking daughters should face a differentially greater mortality risk relative to high-ranking offspring or low-ranking sons (i.e. offspring survival will be predicted by the interaction between offspring sex and maternal rank (Clark 1978, Silk 1983, 1984, van Schaik & Hrdy 1991)).

Finally, the local resource enhancement hypothesis argues that mothers in cooperatively-breeding species will benefit from over-producing whichever sex is better at providing help to developing offspring (Gowaty and Lennartz 1985, Emlen et al 1986, Pen and Weissing 2000). In species with sex-biased dispersal, this would generally be the non-dispersing sex, and results from cooperatively-breeding primates support this prediction (Silk and Brown 2008). Baboons are not cooperative breeders, but they do engage in between-group competition, such that individuals benefit from being in larger groups, up to a point (Markham et al. 2012). At the same time, however, living in groups that are too large results in increased within-group competition (Altmann & Alberts 2003, Beehner et al. 2006, Charpentier et al. 2008, Lea et al. 2015). Because of the conflicting benefits and costs of large group size, the optimal group size appears to be intermediate (Markham et al. 2015).

Combining the insights of the local resource competition and local resource enhancement hypotheses leads to the prediction that female cercopithecine primates will benefit from over-producing philopatric daughters when they are in small, fast-growing groups (causing the groups to grow and attracting more immigrant adult males) and from over-producing dispersing sons when they are in large, slow-growing groups (causing the groups to shrink, or grow more slowly, Table 1). Thus, when the nature of competition is variable over time, the local resource competition and enhancement hypotheses represent two sides of the same coin. Furthermore, group-level sex biases may result from individual-level sex biases that are in line with the predictions of the Trivers-Willard or female rank enhancement hypotheses, such that the relationship between maternal rank and offspring sex might depend on the intensity of competition and vary over time (van Schaik & Hrdy 1991).

### Goals of the current analysis

These four hypotheses produce a combination of overlapping and conflicting predictions about the ways in which offspring sex and survival should be biased (Table 1). The goal of this analysis is to systematically assess each of these alternative predictions using 50 years of data from the Amboseli baboon population in southern Kenya. Using the largest sample size of wild primates available in a single population, we assess whether (1) offspring sex is related to maternal rank, (2) whether this relationship varies over time, and (3) whether female baboons adaptively modulate offspring sex to match the environmental conditions that offspring will experience. In addition to maternal rank, we also consider whether offspring sex and survival are predicted by other indicators of maternal condition, such as exposure to early life adversity (e.g. experiencing drought or maternal loss in early life). We also assess whether the predictions of the local resource competition and local resource enhancement hypotheses hold in this population, by assessing (4) whether mothers bias their offspring towards the dispersing sex, (5) whether such a bias is predicted by measures of competitive intensity (i.e., group size or population growth rate) or their interaction with maternal rank, and (6) whether female offspring are at a differentially increased risk of immature death when born to low-ranking mothers. We fail to find evidence for any of these mechanisms, indicating a lack of any adaptive biasing of offspring sex in this population.

## Methods

### Study population

The Amboseli Baboon Research Project (ABRP) is a long-term, longitudinal study of non-provisioned, individually recognized, wild baboons living in and around Amboseli National Park, Kenya. Demographic, behavioral, and environmental data have been collected on a near-daily basis since the inception of the project in 1971. Critical for the analyses presented here, ABRP has data on offspring conception, birth, and death dates as well as data on female dominance rank (see below) from 1971 to 2020. Additional description of the study population and its history can be found elsewhere (Alberts & Altmann 2012).

### Calculating female social dominance ranks

A detailed description of dominance rank calculations can be found in Hausfater (1975) and Alberts et al. (2003). Briefly, sex-specific dominance ranks are calculated on a monthly basis for all adult males and females relative to other individuals of the same sex in the same social group. Ranks are calculated by generating an NxN matrix (where N is the number of individuals in the social group) that contains symmetrical rows and columns, each corresponding to an individual animal identity. The cells of the matrix contain the number of times that the animal represented by a given row won an agonistic interaction against the animal represented by a given column in that month. The columns and rows of the matrix are then ordered to minimize the number of wins that appear below the diagonal of the matrix. The resulting order of the columns is the ordinal rank (e.g. 1, 2, 3, etc.) of the animals represented by those columns. To calculate proportional rank (the rank metric used in all analyses below), we determine the proportion of other same-sex adults in the group that an individual in question dominates (Levy and Zipple et al. 2020). For example, a female ranked 3 in a group that contains five adult females has a proportional rank of 0.5 (she outranks 2 of the other four females in the group). For the below analyses, we calculated maternal dominance ranks at the time of conception for all live births of known-sex offspring, in all cases where reliable agonism data were available for the mother and other females in her group (n = 333 mothers at the time of n = 1372 live births; reliable dominance ranks are not available for some periods and some groups, e.g., during group fissions).

### Estimating conception dates

When female yellow and anubis baboons (the two species ancestries represented in Amboseli) become pregnant, the skin adjacent to their callosities changes from black to pink (Altmann 1973). By using this pregnancy sign, in combination with sexual swelling and mating data, we are able to identify conception dates within a few days’ precision. This visual method of pregnancy identification has been verified using sex steroid hormone data (Beehner et al. 2006, Gesquiere et al. 2007).

### Estimating annual social group size and growth rates

Using near-daily data on group censuses, we estimated group size and group growth rate for each social group in each year in our dataset. To do so, we calculated the proportional change in the mean number of individuals present in a social group from the year of birth to the year that followed. For example, Alto’s group contained an average of 43.4 individuals on any given day in 1980. In 1981, Alto’s group contained an average of 45.8 individuals. We therefore estimated the growth rate in Alto’s group in 1980 to be 0.055 (2.4/43.4).

### Testing the Trivers-Willard and female rank enhancement hypotheses across all social groups and years

Although many studies of rank-related sex biases consider the proportion of sons born to females of a given rank, here we instead consider the proportion of daughters born (a statistically equivalent approach). We do so because hypotheses about baboon sex bias are generally related to mothers’ ability to influence the rank of their daughters, but not their sons.

We tested for a relationship between maternal rank at the time of conception and offspring sex at two different scales. First, we used data from all live births for which relevant rank data were available from across the entire study (n = 1372 live-born offspring) to build a mixed effects logistic regression model (R package: ‘glmmTMB’) that predicted the sex of each live-born offspring as a function of its mother’s rank, with maternal identity included as a random effect (Magnusson et al. 2017).

Second, we used the same analytical approach to ask whether there were some periods or social groups in the history of the ABRP when offspring sex was significantly predicted by maternal rank. We already knew that one such period existed—Altmann (1980) and Altmann et al. (1988) had previously described the strong relationship between maternal rank and offspring sex during the first decade of observation of Alto’s group, a social group observed between 1971 and its permanent fission in 1992. To test whether other social groups showed the same pattern during some periods, we built a series of mixed effects logistic regression models. Each model was built from data collected from a single social group over a seven-year period. In each model, the response variable was the sex of each offspring born during that seven-year period, and the predictor variable was maternal dominance rank; maternal identity was included in each model as a random effect.

We chose a seven-year period because it is similar to the original time window analyzed by Altmann (1980), which demonstrated a statistically significant bias in offspring sex ratio as a function of maternal rank. Thus, we reasoned that considering data from seven years in a single group effectively balanced the analytical benefits of increased sample size against the costs of increased environmental heterogeneity during longer periods of observation (see introduction). Using a time period comparable to Altmann (1980) also allowed us to perform an analytical thought experiment in which we asked whether the previously published relationship would have been identified if data collection had started in a different social group at a different time. We considered only subsets of data from groups for which seven consecutive years of birth data could be analyzed. For example, Alto’s group (Group 1) fissioned into two social groups in 1990. The latest subset of data considered from Alto’s group therefore spanned 1984-1990. We included only 7-year periods with at least 10 births recorded during that period, resulting in a total of 109 overlapping periods across all study groups.

### Testing the Trivers-Willard and female rank enhancement hypotheses: do females adaptively modulate the direction and magnitude of a rank-related offspring sex bias?

Daughters born to high-ranking mothers may be advantaged relative to sons under some conditions, but disadvantaged under other conditions. If so, we predict temporal and between-group variability in sex-ratio biasing as a function of temporal variability in the survival of daughters born to high-ranking mothers.

To test this possibility, we used a 3-step analysis. The first step was to build Cox proportional hazards models of offspring survival (hereafter ‘survival models’) for the first four years of life. This is just prior to puberty for most females (who achieve menarche at a median age of 4.5 years in Amboseli) and around the earliest age of dispersal for males (median dispersal age is 7.6 years; median age at testicular enlargement for males is 5.4 years) (Onyango et al. 2013). We modeled offspring survival as a function of maternal rank, offspring sex, and the interaction between maternal rank and offspring sex (R package: ‘survival’, (Therneau & Lumley 2015)). The magnitude of this interaction term is the measure of interest—we want to know whether, in a given period, the difference in survival between daughters and sons was greater for high-ranking than for low-ranking mothers. If such rank-related differences in the survival of daughters and sons exist, and if they vary across time and social groups, then mothers could theoretically benefit from modulating the magnitude of a rank-related sex bias to mirror variation in sex-associated survival differences.

To assess whether temporal variability affects sex-ratio biasing, we ran our survival models used data from multiple non-overlapping periods of time for each social group. Unlike the overlapping time window analysis above, we used non-overlapping windows in this case to enforce greater independence between analyses. The lengths of these non-overlapping time periods varied: because different groups were under study for different lengths of time we were unable to split groups’ periods of observations into equally sized lengths. For groups that existed in our dataset for 10 years or less, we used the entire period of observation of that group as an independent unit of analysis. For groups that existed for more than 10 years, we split their contribution into two approximately equal time windows and treated the two windows as separate units of analysis. In combination, these two procedures produced 18 separate group-time windows that were a minimum of 6 years and a maximum of 10 years long. For example, the data from Alto’s group (1971-1990) could readily be split into two, ten-year subsets (1971-1980, 1981-1990). In contrast, because of the relatively short observation time for Acacia’s group, we retained all of the data from this group (2013-2020) as a single, eight-year set of data.

Because of the importance of knowing infants’ ages with precision, we included in this analysis only those infants whose birth date was known within a few days’ error. We also excluded from the survival analysis (but not from other analyses) any offspring born into non-wild feeding groups, which gain a substantial portion of their daily caloric intake from human food waste, as well as any infants born into groups with less than 6 years of total data. Our final sample for the survival analyses contained 1121 live-born offspring.

After calculating temporal and between-group variability in rank-related differences in survival between male and female offspring, we then calculated the magnitude of the effect of maternal rank on offspring sex for the same non-overlapping time windows. We calculated coefficient estimates using the same mixed-effects logistic regression models described above (“*Testing the Trivers-Willard and female rank enhancement hypotheses across all social groups and years*”). Together, these two analyses yielded estimates of (i) variability in the potential benefits that mothers could accrue if they biased their offspring’s sex in the proper direction to maximize survival and (ii) variability in the estimated association between maternal rank and offspring sex across groups and time periods.

The third step in our analysis was to test the prediction that these two measures are related to each other in the manner predicted by adaptive hypotheses for sex-ratio biasing. If females adaptively modulate the direction and magnitude of a rank-related sex bias in their offspring, we expect a significant positive relationship between the effect of maternal rank on offspring sex and the interaction effect between offspring sex and maternal rank on offspring survival. In other words, mothers of different ranks should bias their offspring production towards the *right sex*, under the *right conditions*. To test this prediction, we built a linear regression that predicted the coefficient of the rank terms from the logistic regression models (the result of step two) as a function of the interaction terms from the survival models (the result of step one).

### Are the Trivers-Willard or female rank enhancement hypotheses supported by considering other measures of maternal condition?

Inspired by the previous literature, the analyses described above focus on rank as the primary indicator of condition. However, it is also possible that females alter their sex ratio in response to their physical condition, but that maternal rank is a poor proxy. To test this possibility, we assessed whether alternative measures of maternal condition predict offspring sex ratio. First, we used mixed effects logistic regression models to ask whether offspring sex was predicted by whether mothers experienced each of five sources of early life physical and social adversity prior to maturity, and/or a cumulative measure of these five adverse experiences (early drought, high group density, maternal loss, low maternal rank, presence of a close-in-age younger sibling; see Tung and Archie et al 2016 for a description of sources of adversity). Experiencing early life adversity is associated with dramatically shorter lifespans for female baboons in the Amboseli population (Tung and Archie et al. 2016) as well as reduced offspring survival (Zipple et al. 2019, 2021). Because the early adversity analysis required data on the early-life conditions faced by mothers, the sample size for this analysis was substantially lower than for other analyses (n = 742 live births for which data were available on five sources of early-life adversity for the mother, versus n = 1372 for the preceding analyses).

The second alternative measure of condition that we considered was whether females were near the end of their lives, as indicated by their death within 1, 2, or 4 years of their offspring’s birth (analyzed as three separate models). We expected mothers to be in worse condition in the years before they died, and that offspring would survive less well when their mothers were within a few years of death (Zipple et al. 2019, 2021). The three maternal survival analyses required data on whether the mother in question survived a given period following offspring birth, which reduced our sample size in these analyses to varying degrees (n = 1214 live births for 4-year analysis, 1301 for 2-year analysis, and 1343 for 1-year analysis).

### Testing the local resource competition and enhancement hypotheses

Lastly, to test whether females in Amboseli exhibit a global bias towards producing males (the dispersing sex; see Local Resource Competition, Table 1), we performed a two-sided, two-proportions z-test (R function ‘prop.test’) using all offspring in the dataset. To assess context/environment-specific predictions of the local resource competition and enhancement hypotheses, we built mixed logistic regression models that predicted offspring sex as a function of either group size or population growth rate (two measures of intensity of competition). We also tested whether either of these measures of competition significantly interacted with maternal rank to predict offspring sex. Finally, we asked whether there was a significant interaction between maternal rank and offspring sex in predicting offspring survival to 4 years of age (the approximate age of female maturation and the earliest age of male dispersal; Onyango et al. 2013). We did not calculate social group size estimates for offspring born in the year that groups fissioned, fused, or were dropped from observation, nor did we calculate growth rate estimates for offspring born in the year of or the year prior to such an event. As a result, sample size was reduced for analyses involving group size (n = 1274 live births) or population growth rate estimates (n = 1109 live births).

## Results

### Testing the Trivers-Willard and female rank enhancement hypotheses across all social groups and years

In the pooled, 50-year dataset, maternal rank at the time of conception did not predict offspring sex (estimate from mixed effects logistic regression = 0.12, se = 0.18, z = 0.68, p = 0.50, n =1372 live births). A visual inspection of the data further reinforces that there is no consistent relationship between an offspring’s sex and the rank of its mother (Figure 1).

**Figure 1.**
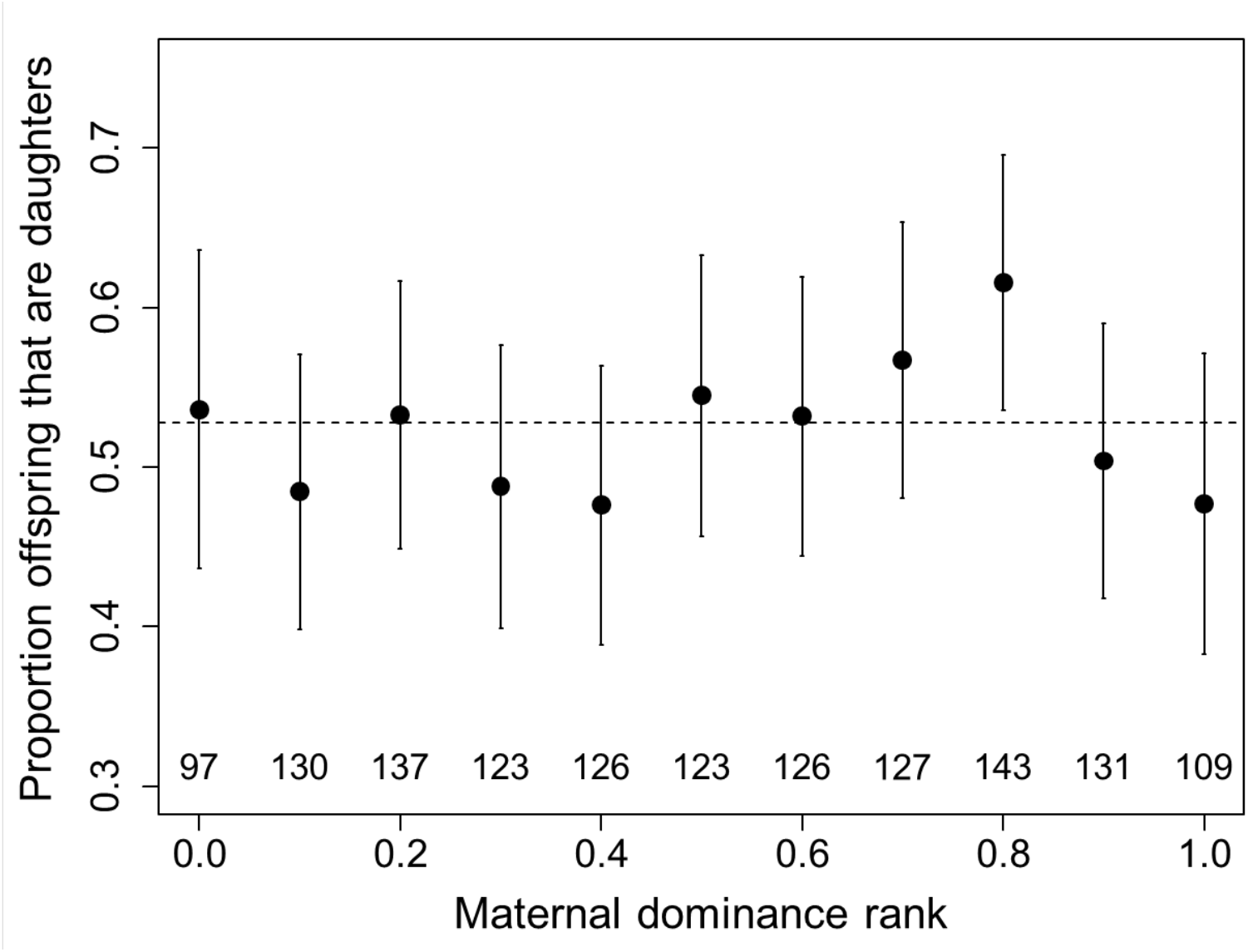
The proportion of daughters born to mothers falling into each decile of maternal dominance rank for 1372 live births. Error bars indicate 95% confidence intervals for each decile, the dashed line indicates the population mean proportion of daughters (52%), and the numbers below each point indicate the number of offspring included in each point. Although displayed as bins in this illustrative figure, proportional ranks were treated continuously in the logistic regression model that we report in the main text, in which we found no relationship between maternal rank and offspring sex (p = 0.50).

In addition to the pooled analysis, we also tested whether maternal rank predicted offspring sex in some groups and some years. In total, we assessed whether offspring sex was significantly predicted by maternal rank in 109 subsets of the data representing successive overlapping 7-year time spans in single social groups. During some periods in some groups, high-ranking mothers had far more daughters than low-ranking mothers, while in other periods and groups the trend was reversed. Some of this variation is due to variation in sample size for analyses in different periods, which ranged from 11 to 100 offspring (subsets containing 10 or fewer infant births were excluded). Periods with smaller sample sizes generally had larger absolute effect size estimates, consistent with well-known winner’s curse effects (see supplemental Figure S1). Overall, this analysis yielded four major results.

First, we identified enormous variation in the estimated magnitude of the association between maternal rank and offspring sex in different groups at different times. However, in the vast majority of time periods, maternal rank did not predict offspring sex at a statistically significant level (alpha = 0.05). Maternal rank significantly predict offspring sex in 6% (6/109, p < 0.05) of all 7-year time periods, but none of these results survive a Bonferroni correction for multiple hypothesis testing.

Second, the magnitude of the maternal-rank effect during the first 10 years of the study (as indicated by the first four points on the dark red line of Figure 2 [Alto’s group]) was large and statistically significant at a nominal p-value of 0.05, corresponding to a scenario in which an offspring had an ∼88% chance of being female if born to the highest-ranking female in the group and only an ∼18% chance of being female if born to the lowest-ranking female in the group.

**Figure 2.**
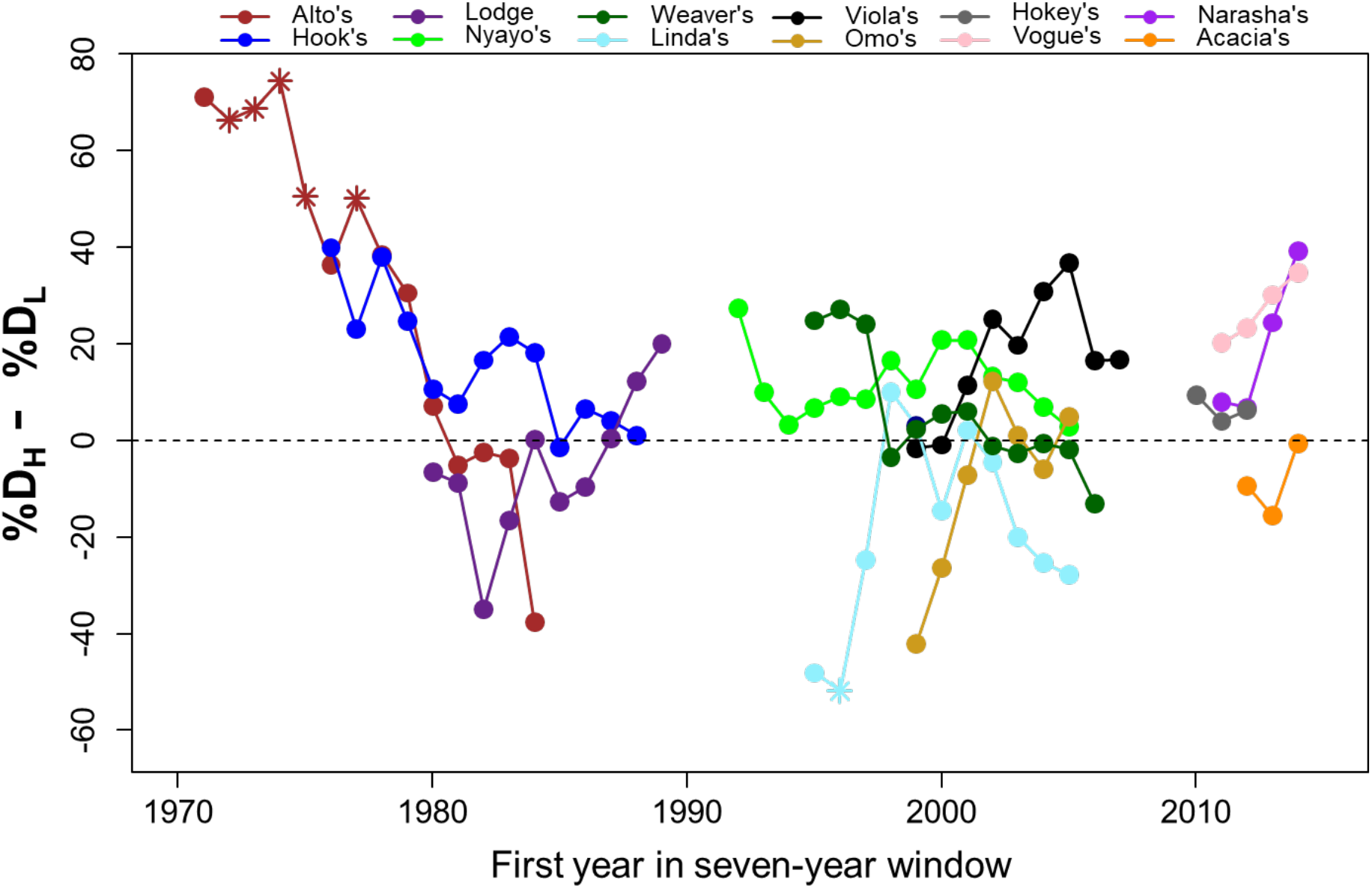
The magnitude of the estimated difference in the percentage of female offspring born to the absolutely highest-ranking mothers (%DH) and lowest-ranking mothers (%DL) in different groups over different time periods. Each point represents a unique estimate extracted from a logistic regression model of offspring sex as predicted by maternal proportional rank, using data from a single group over a seven-year period that begins at the x-value of that point (e.g., an x-value of 1971 represents data in a single group from 1971-1977). Asterisks indicate groups and times in which models indicated that maternal rank significantly predicted offspring sex (p < 0.05), with group identities indicated by different colors and the dashed line indicating the null expectation. For example, to calculate the y-value of the first red point, we built a mixed effects logistic regression model of offspring sex as predicted by maternal rank, using data from all offspring born between 1971 and 1977 in Alto’s group (inclusive, representing an x-value of 1971). We then used the resulting model output to estimate the difference in the percentage of daughters born to the highest-ranking mothers (91%) and the percentage of daughters born to the lowest ranking-mothers (20%) from 1971-1977, yielding a point with a y-value of 71%.

These points correspond to the striking observation captured by the original analyses in Altmann (1980) and Altmann et al. (1988).

Third, this early period of observation in Alto’s group appears quite atypical compared to all time periods in other study groups as well as other time periods in Alto’s group. Excluding these first four seven-year time windows for Alto’s group, the median estimated difference in the proportion of female offspring born to the highest and lowest ranking females in each group-time window combination was only 0.05 across the study period, as compared to an average difference of 0.70 in these first four subsets.

Fourth, early in the group history for Linda’s group (dark blue line beginning in 1995), there was a similarly large estimated difference between the proportion of daughters born to the highest- and lowest-ranking females, but in the opposite direction of the difference observed early in the observation period for Alto’s group. Specifically, only ∼14% of the offspring of the highest-ranking females were expected to be daughters, as compared to ∼66% of the offspring of the lowest-ranking females.

In sum, we did not find any evidence that females consistently exhibit a rank-dependent strategy of biasing offspring sex ratio (Figure 2). If long-term data collection from our population had started at essentially any other time or in any other social group, researchers would not have identified an apparent relationship between maternal rank and offspring sex, with the exception of Linda’s in the mid-1990s, when a significant effect would have appeared in the opposite direction.

#### Testing the Trivers-Willard and female rank enhancement hypotheses: do females adaptively modulate the direction and magnitude of a rank-related offspring sex bias?

The variation we found in the relationship between maternal rank and offspring sex (Figure 2) might be the result of random processes. Alternatively, it might map onto variation in which offspring sex experienced the highest survival probability, given maternal rank and the environment into which the offspring was born. If mothers adaptively modulate their offspring sex ratios in this way, we would expect to find a positive relationship between the magnitude of any maternal rank-related sex bias in a given period and the magnitude of the survival advantage to offspring of the ‘right’ combination of sex and maternal rank in the same period (see Methods). In other words, under an adaptive scenario we would predict that high-ranking mothers only bias their offspring sex ratio towards daughters *when they generate a survival advantage for their offspring by doing so*.

In contrast to this prediction, the coefficient estimates for sex-biased survival and maternal rank-related sex bias were not correlated. Specifically, the magnitude of the relationship between maternal rank and offspring sex in a given period did not predict the magnitude of the interaction between maternal rank and offspring sex on offspring survival in the same period (R^2^ = 0.08, p = 0.25 Figure 3). Additionally, only 6 out of 18 periods fell in the first and third quadrants of Figure 3, which are consistent with maternal rank-dependent adaptive modulation of offspring sex ratio (the bottom left and upper right quadrants of Figure 3). In contrast, 12 of the time periods we considered fell in quadrants two and four, which are associated with *costly* modulation of rank related effects (i.e., high-ranking mothers producing offspring of the “wrong” sex). Thus, this analysis provides no evidence that females adaptively change the magnitude or direction of a rank-related sex bias in order to maximize the survival prospects of their offspring.

**Figure 3.**
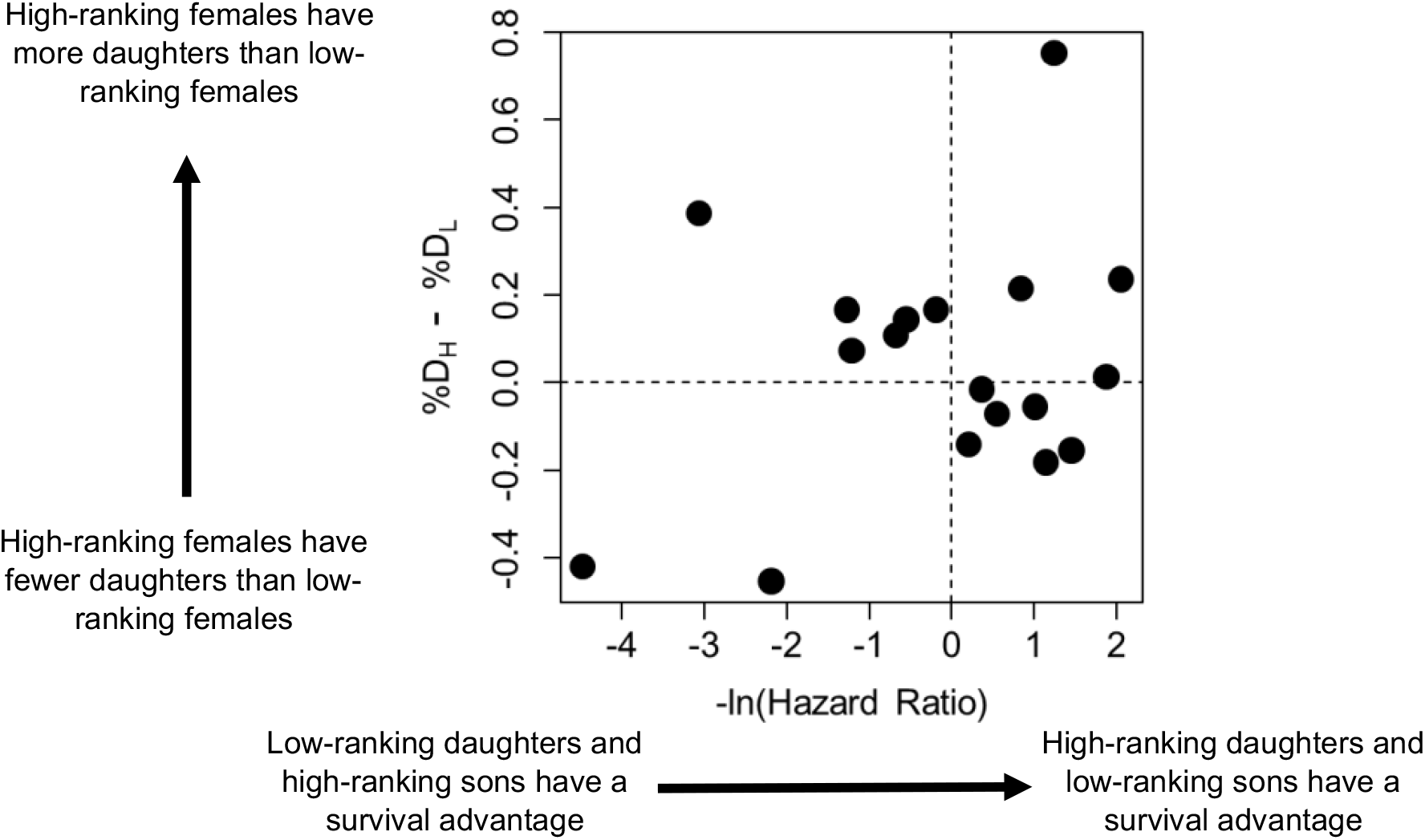
The magnitude of the survival benefit experienced by daughters born to high-ranking mother (x-axis) does not predict the magnitude of the maternal rank-based bias in offspring sex (y-axis). Each of the 18 points represents model estimates from six to ten year time windows for a single social group. The x-axis represents the coefficient of the interaction between maternal rank and offspring sex in a Cox proportional hazards model of offspring survival (see Methods for additional details). Positive values on the x-axis indicate periods when offspring benefitted from being female when born to high-ranking mothers or from being male when born to low-ranking mothers. The y-axis represents the magnitude of the rank-related sex bias during a given time period (see Figure 2 for a detailed description of %DH - %DL). Positive values on the y-axis indicate periods when high-ranking females had more daughters than low-ranking females (%DH > %DL, see Figure 2). Adaptive modulation of offspring sex ratio to maximize offspring survival would predict a positive association between these two estimates, but neither linear regression (to test for an overall correlation: R^2^ = 0.08, p = 0.25) nor a Fisher’s Exact Test (to test for directional concordance: p = 0.19, in opposite direction as predicted) identifies such an association.

#### Are the Trivers-Willard or female rank enhancement hypotheses supported by considering other measures of maternal condition?

Although maternal early adversity strongly predicts offspring survival (Zipple et al 2019, 2021), maternal early life adversity does not predict offspring sex in either multivariate or cumulative adversity models (Table 2). Similarly, although impending maternal death predicts lower offspring survival, maternal death in the one-, two-, or four-year periods following offspring birth also did not predict offspring sex (Table 2).

**Table 2.** Model results from mixed effects logistic regression models that predict offspring sex as a response to alternative measures of maternal condition.

| *Measure of Female Condition* | *Coefficient Estimate<sup>^</sup>* | *Estimated Effect on proportion of female offspring* | *p value* |
| --- | --- | --- | --- |
| ***Early Life Adversity (n = 807)*** |  |  |  |
| *Multivariate Model* |  |  |  |
| Early Maternal Loss | -0.17 | -0.04 | 0.34 |
| Close-In-Age Younger Sibling | -0.05 | -0.01 | 0.77 |
| Born to a Low-Ranking Mother | 0.12 | 0.03 | 0.51 |
| Born in a Large Group | 0.49 | 0.12 | 0.13 |
| Born During a Drought | 0.41 | 0.10 | 0.13 |
| *Cumulative Adversity Model* |  |  |  |
| Cumulative Adversity | 0.06 | 0.01 | 0.54 |
| ***Impending Maternal Death*** |  |  |  |
| Within 1 Year of Offspring Birth (n = 1343) | 0.13 | 0.03 | 0.56 |
| Within 2 Years of Offspring Birth (n = 1301) | 0.10 | 0.02 | 0.54 |
| Within 4 Years of Offspring Birth (n = 1214) | 0.24 | 0.06 | 0.07 |

<sup>^</sup> Positive values indicate an increase in the proportion of daughters.

#### Testing the local resource competition and enhancement hypotheses

Overall, the global sex ratio at birth for the Amboseli baboon population did not significantly deviate from parity (52% females, 95% CI = <0.50 – 0.55, p = 0.07). Notably, even the suggestion of a deviation from parity—in favor of females—is in the opposite direction to that predicted by the local resource competition hypothesis, which predicts a bias towards males, the dispersing sex. Offspring sex was not predicted by group size in the year of birth (p = 0.10, coef. estimate = - 0.005), or by social group growth rate (p = 0.35, coef. estimate = -0.74) in bivariate models predicting offspring sex along with a random effect of maternal ID. In more complex models (Table 3), offspring sex was also not predicted by the interaction between maternal rank and either group size in the year of birth (p = 0.88) or social group growth rate (p = 0.26, Table 3). Overall, offspring survival was not significantly predicted by the interaction between maternal rank and offspring sex (p = 0.93), though the main effect of maternal proportional rank was a significant predictor of offspring survival (HR = 0.40-0.99, p < 0.05), consistent with previous results in this population (Silk et al 2003, Zipple et al. 2019). See Table 3 for full results.

**Table 3.** Model results from mixed effects models that predict offspring sex and offspring survival as a response to alternative measures of competitive environment

| *Model/Parameter\** | *Coefficient*<br>*Estimate<sup>^</sup>* | *Std. Error* | *p Value* |
| --- | --- | --- | --- |
| ***Offspring Sex ~ Group Size (n = 1274)*** |  |  |  |
| Maternal Proportional Rank | 0.28 | 0.57 | 0.63 |
| Group Size in Year of Birth | -0.004 | 0.006 | 0.48 |
| Mat. Rank x Group Size | -0.001 | 0.009 | 0.88 |
| ***Offspring Sex ~ Group Growth Rate (n = 1109)*** |  |  |  |
| Maternal Proportional Rank | 0.41 | 0.24 | 0.09 |
| Social Group Growth Rate | 0.67 | 1.45 | 0.65 |
| Mat. Rank x Growth Rate | -2.74 | 2.44 | 0.26 |
| ***Offspring Survival ~ Sex and Rank (n = 1242)*** |  |  |  |
| Maternal Proportional Rank | -0.46 | 0.23 | **0.046** |
| Offspring Sex | 0.07 | 0.18 | 0.77 |
| Mat. Rank x Offspring Sex | 0.03 | 0.31 | 0.83 |

\*All models include maternal identity as a random effect.
<sup>^</sup>Positive values indicate an increase in the proportion of daughters (offspring sex models) or an increase in mortality (survival model)

## Discussion

Overall, we find no evidence that female baboons adaptively bias the sex ratio of their offspring in the direction predicted by the competing hypotheses in Table 1. First, we find no evidence that female baboons consistently alter the sex ratio of their offspring based on their dominance rank or other metrics of female condition (Figure 1, Table 2). While it is difficult to rule out the possibility that females engage in such rank-based biasing in some very specific environmental contexts, our data demonstrate that such a strategy is, at best, quite rarely employed (Figure 2). Next, we find no evidence that females modulate their offspring’s secondary sex ratios in favor of the sex that is more likely to survive based on the females’ social rank and the environmental conditions that the offspring experiences (Figure 3). Specifically, while we acknowledge that this particular analysis relies on point estimates from model outputs with substantial uncertainty, the lack of any evidence for an effect means that we can be confident that any relationship between the magnitude of the survival benefits of a rank-related sex bias and the rank-related sex bias that actually exists is, at best, very weak. Finally, we find no relationship between offspring sex ratio and measures of competitive intensity, nor do we find support for an interaction between competitive intensity and maternal dominance rank (Table 3). While the predictions of each hypothesis in Table 1 differ, they all predict that offspring sex ratio or offspring survival will be shaped by some combination of maternal rank, group size, and population growth rate. No such relationship is detectable in our dataset, which represents the largest sample size (n =1372) from a single wild primate population ever used to assess these hypotheses.

The absence of a relationship between maternal rank and offspring sex in this study contradicts previously published results from our study system based on data from the beginning of long-term observations, from 1971-1981 (Altmann 1980, Altmann et al. 1988). Those results are reproduced in our analysis of the data from that period, and indeed the strength of the relationship between maternal rank and offspring sex in that data set is striking (see Figure 2, above and Figure 25.3 in Altmann et al. 1988). The fact that other time periods in the Amboseli baboon data set, and other primate populations, do not show a similar pattern, suggests that this previous result was a false positive (Type I error) (Brown & Silk 2002, Silk et al. 2005). If long-term data collection on the Amboseli baboons had started during essentially any other time period or in any other social group, researchers would not have identified an apparent relationship between maternal rank and offspring sex.

At the same time, the rank-related results from this analysis are consistent with previous theoretical work by Altmann and Altmann (1991). Altmann and Altmann (1991) modeled the group-level demographic implications of rank-related modulation of offspring sex ratios in a matrilocal species in which females inherit their mother’s dominance rank (such as baboons and some other cercopithecine monkeys). They showed that if high-ranking females in such a species biased their offspring sex ratio towards sons, the result would be an unstable group size, such that small groups rapidly decline in size and large groups grow at an ever-increasing rate (Altmann & Altmann 1991). In contrast, if high-ranking females were to bias their offspring sex ratio towards daughters, then group size would be highly regulated: groups would remain stable at a near constant size, composed primarily of closely related females (Altmann & Altmann 1991). Neither of these outcomes is consistent with the empirical dynamics of baboon social groups where, when population growth is positive overall, small social groups grow in size and large social groups continue to grow until they fission into smaller groups (Van Horn et al. 2007, Markham et al. 2015). Further, rather than being tightly regulated around a stable group size, group sizes vary widely in the Amboseli population from less than 10 to more than 100 individuals (Stacey 1986, Markham et al. 2015). Thus, the dynamics of social group sizes in the Amboseli baboons are counter to the demographic predictions that ensue from maternal manipulation of offspring sex ratio, providing a separate line of evidence that such manipulation does not happen in this population.

One possible explanation for the apparent absence of a relationship between maternal rank and offspring sex in our population is that female rank may not be a good proxy of female “condition” in nonhuman primates. This possibility is unlikely to explain the results from our population for two reasons. First, female rank has been associated with a wide range of traits that are likely to be related to condition in our population, including offspring survival, inter-birth interval, attainment of sexual maturity, and the strength of social relationships (Silk et al. 2003, Charpentier et al. 2008, Archie et al. 2014, Gesquiere et al. 2018, Zipple et al. 2019, Levy et al. 2020). Second, we also fail to observe a relationship between other metrics of maternal condition and offspring sex. Specifically, offspring sex is not predicted by maternal early life adversity nor by their mother’s survival in the earliest years following their birth (Table 2), both of which predict offspring survival overall and likely reflect maternal condition (Tung and Archie et al. 2016, Zipple et al. 2019).

Why have females failed to evolve the ability to manipulate the secondary sex-ratio of their offspring to their benefit, as predicted by the female rank enhancement and local resource competition hypotheses? We documented an enormous range in the interaction between offspring sex and maternal rank on offspring survival (Figure 3), indicating that females *could* derive substantial benefit by producing offspring of the ‘right’ sex at any given time, depending on the survival prospects of the offspring. At least two barriers may prevent the evolution of such a strategy.

First, the mechanisms available to female mammals for influencing offspring sex remain mostly theoretical. Further, even these theoretical mechanisms would operate in the direction opposite to that predicted by the female rank enhancement and local resource competition hypotheses (as applied to baboons), which predict that good-condition females should produce more daughters and poor-condition females should produce more sons (reviewed in (Douhard 2017)). For example, one proposed mechanism is based on the idea that higher-ranking females produce higher levels of circulating testosterone, which could potentially make their oocytes more receptive to Y-chromosome sperm (Grant & Chamley 2010, Douhard 2017). Another possible mechanism suggests that high levels of glucose (consistent with good maternal condition) may lead to higher levels of female embryonic mortality and support male embryonic development (Cameron 2004, Douhard 2017). Finally, some have speculated that high levels of glucocorticoid concentrations (consistent with poor-condition mothers) could lead to differential male embryonic mortality (Navara 2010, Douhard 2017). Each of these potential mechanisms is consistent with the Trivers-Willard hypothesis, and in red deer and bighorn sheep, good maternal condition has been reported to predict the production of sons, although only in specific circumstances (Kruuk et al. 1999, Douhard et al. 2016). However, no mechanisms have yet been proposed through which good-condition mothers could bias their offspring sex ratio towards daughters, but poor-condition mothers the reverse.

Second, even if mechanisms exist that would allow females to facultatively adjust the sex ratio of their offspring, the ability to do so adaptively relies on females’ ability to use cues available at the time of conception to identify the fitness-favoring sex in *future* environmental conditions. In the case of baboons (a relatively long-lived mammal), this would require females to accurately assess whether environmental conditions over the coming years and decades would differentially benefit the reproductive success of male versus female offspring, given her social rank and environmental cues at the time of conception. In the highly dynamic physical and social environment that female baboons experience in the Amboseli population, such an assessment is likely to be impossible. Thus, the results presented here add to a growing body of evidence from this and other populations that early life environmental cues may not be sufficiently informative to select for predictive adaptive responses that optimally align with future environmental conditions (Hayward & Lummaa 2013, Douhard et al. 2014, Lea et al. 2015, Weibel et al. 2020)

This inability to predict the future may also explain the absence of any relationship between group size or group growth rate and offspring sex. Although it may be beneficial to females to modulate their offspring’s sex depending on future group size, females likely lack sufficiently reliable information to make such a determination at the time of conception. Small groups tend to grow faster than large groups and large groups tend to fission, but this is a very noisy process that proceeds quite differently in different social groups (Stacey 1986, Markham et al. 2015). As a result, any individual female is unlikely to be able to predict future group size or competitive environment based on group size or growth rate at the time of conception. Importantly, mistakes would be costly, as differential death or abortion of a fetus in a singular breeder like baboons can have a meaningful effect on lifetime reproductive success. Further, even the benefits of making a “correct decision” may be less than they appear: a mother that aborted a fetus of the disadvantageous sex would have only an ∼50% chance of conceiving an offspring of the advantageous sex the next time she became pregnant, so selectively aborting a fetus of the wrong sex (as required by all proposed mechanisms above) would substantially slow female reproductive life histories.

In sum, evolutionary hypotheses about facultative adjustment of offspring sex ratio are compelling. Yet, among primates there remains no convincing evidence that condition-dependent manipulation of offspring sex systematically occurs. The sum of the evidence from more than a dozen species instead indicates that offspring sex is independent of maternal condition. Thus, between-species variation in secondary sex ratio certainly exists, but variation within primate species does not appear to depend on maternal condition—or at least not strongly enough that it is dependably detectable even in the best-powered analyses to date (Brown & Silk 2002, Silk et al. 2005, Silk & Brown 2008). On the other hand, the Trivers-Willard hypothesis has been partially supported in at least two ungulates: red deer and big-horn sheep (Clutton-Brock et al. 1984, Kruuk et al. 1999, Douhard et al. 2016).

It may be that observations reported in ungulates reflect historical false positives similar to that which we report here. But if not, the apparent difference between primates and ungulates motivates a central question to be addressed going forward: what explains why offspring sex in (some) ungulates appears to be dependent on maternal condition, while the same does not appear to be true in primates? One possible explanation is that the fitness of sons is more tightly tied to maternal condition in ungulates than in primates (see Altmann 1980). This possibility could be tested by identifying those exceptions that prove the rule in both taxa. That is, if there are male primates whose fitness outcomes depend on maternal condition, these are the species in which we would be most likely to see sex ratios dependent on maternal condition. Conversely, if there are ungulate species in which male fitness outcomes are independent of maternal condition, we would expect these to be species in which offspring sex would also be independent of maternal condition. The first step towards such a test is a more complete assessment of the presence or absence of a relationship between maternal condition and offspring sex in more populations of wild mammals, as the number of species for which we can have confidence in this assessment remains low (Brown 2001, Brown & Silk 2002, Silk et al. 2005).

## Acknowledgments

MNZ is supported by a Klarman Postdoctoral fellowship at Cornell University as well as an NSF Postdoctoral Research Fellowship in Biology (grant # 2109636). We gratefully acknowledge the support of the National Science Foundation and the National Institutes of Health for the majority of the data represented here, currently through NSF IOS 1456832, and through NIH R01AG053308, R01AG053330, R01AG071684, R01HD088558, R01AG075914, and P01AG031719. We also thank Duke University, Princeton University, and the University of Notre Dame for financial and logistical support. In Kenya, our research was approved by the Wildlife Research Training Institute (WRTI), Kenya Wildlife Service (KWS), the National Commission for Science, Technology, and Innovation (NACOSTI), and the National Environment Management Authority (NEMA). We also thank the University of Nairobi, the Institute of Primate Research (IPR), the National Museums of Kenya, the members of the Amboseli-Longido pastoralist communities, the Enduimet Wildlife Management Area, Ker & Downey Safaris, Air Kenya, and Safarilink for their cooperation and assistance in the field. Particular thanks go to S. Sayialel for data collection in the field, and to T. Wango and V. Oudu for their untiring assistance in Nairobi. The baboon project database, Babase, is expertly managed by N. Learn and J. Gordon. Database design and programming are provided by K. Pinc. This research was approved by the IACUC at Duke University, University of Notre Dame, and Princeton University and adhered to all the laws and guidelines of Kenya. For a complete set of acknowledgments of funding sources, logistical assistance, and data collection and management, please visit http://amboselibaboons.nd.edu/acknowledgements/.

## Supplemental Material

**Figure S1.**
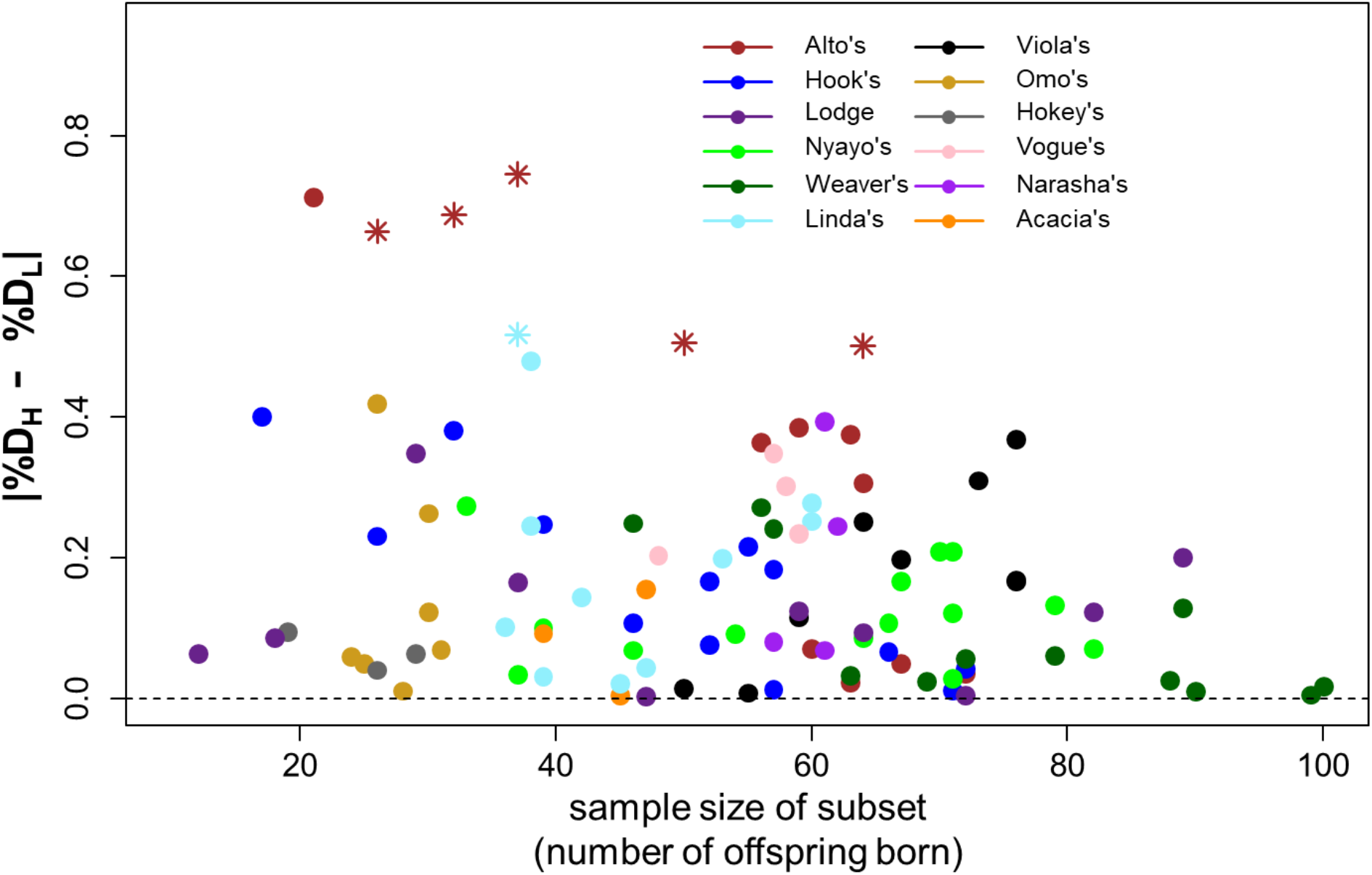
The largest apparent effects of maternal rank on offspring sex occurred during analyses of 7-year subsets of the data characterized by smaller sample sizes of offspring born. Plotted here are the absolute values of the estimated magnitude of the sex bias effects previously shown in Figure 2 (the y-axis) versus the number of offspring born during the 7-year group-period contained in each analysis (the x-axis). As in Figure 2, asterisks indicate periods and groups in which models indicated that maternal rank significantly predicted offspring sex, with group identities indicated by different colors, and the dashed line indicating the null expectation. As expected, the average magnitude of the estimated effect declines as sample size increases.

